# Epithelial and neutrophil interactions and coordinated response to *Shigella* in a human intestinal enteroid-neutrophil co-culture model

**DOI:** 10.1101/2020.09.03.281535

**Authors:** Jose M. Lemme-Dumit, Michele Doucet, Nicholas C. Zachos, Marcela F. Pasetti

## Abstract

Polymorphonuclear neutrophils (PMN) are recruited to the gastrointestinal mucosa in response to inflammation, injury, and infection. Herein, we report the development and the characterization of an *ex vivo* tissue co-culture model consisting of human primary intestinal enteroid monolayers and PMN, and a mechanistic interrogation of PMN-epithelial cell interaction and response to *Shigella*, a primary cause of childhood dysentery. Cellular adaptation and tissue integration, barrier function, PMN phenotypic and functional attributes, and innate immune responses were examined. PMN within the enteroid monolayers acquired a distinct activated/migratory phenotype that was influenced by direct epithelial cell contact as well as by molecular signals. Seeded on the basal side of the intestinal monolayer, PMN intercalated within the epithelial cells and moved paracellularly toward the apical side. Co-cultured PMN also increased basal secretion of IL-8. *Shigella* added to the apical surface of the monolayers evoked additional PMN phenotypic adaptations, including increased expression of cell surface markers associated with chemotaxis and cell degranulation (CD47, CD66b, and CD88). Apical *Shigella* infection triggered rapid transmigration of PMN to the luminal side, NET formation as well as bacterial phagocytosis and killing. *Shigella* infection modulated cytokine production in the co-culture; apical MCP-1, TNF-α, and basolateral IL-8 production were downregulated, while basolateral IL-6 secretion was increased. We demonstrated, for the first time, PMN phenotypic adaptation, mobilization, and coordinated epithelial cell-PMN innate response upon *Shigella* infection in the human intestinal environment. The enteroid monolayer-PMN co-culture represents a technical innovation for mechanistic interrogation of gastrointestinal physiology, host-microbe interaction, innate immunity, and evaluation of preventive/therapeutic tools.

**Importance:** Studies of mucosal immunity and microbial host cell interaction have traditionally relied on animal models and *in vitro* tissue culture using immortalized cancer cell lines, which render non-physiological and often unreliable results. Herein we report the development and characterization of an *ex vivo* enteroid-PMN co-culture consisting of normal human intestinal epithelium and a mechanistic interrogation of PMN and epithelial cell interaction and function in the context of *Shigella* infection. We demonstrated tissue-driven phenotypic and functional adaptation of PMN and a coordinated epithelial cell and PMN response to *Shigella*, a primary cause of dysentery in young children in the developing world.

## Introduction

The intestinal epithelium creates a physical and molecular barrier that protects the host from potentially damaging elements in the constantly changing outside environment. Epithelial barrier function is supported by a diverse population of underlying immune cells, which deploy a variety of host-defense mechanisms against harmful agents (1). Coordinated signals resulting from microbial sensing, cell-to-cell contact, cytokines, and other chemical mediators determine the type and extent of responses of gut immune cells, balancing tissue homeostasis with effective anti-microbial function via inflammation.

Advances in understanding intestinal physiology, pathophysiology, and host immunity have traditionally relied on studies conducted in animal models (or animal tissue) and in traditional tissue culture systems using colon cancer cell lines. Animal models, including the use of mutant mouse strains, have contributed to the mechanistic understanding of the composition, function, regulatory processes, and operatives of immunity at the gut mucosa. Unfortunately, host restrictions limit the utility and value of animal models (2, 3). This is the case for many enteric pathogens for which small animals fail to recreate disease as it occurs in humans. Likewise, immortalized (transformed) cell lines (e.g., HT-29, Caco-2, and T84) do not reflect human physiological responses but rather the aberrant behavior of diseased cells (e.g., karyotype defects). These cell-line based cultures also lack the multicellular complexity of the human intestinal epithelium, which further reduces the reliability and significance of the data generated. The establishment of human enteroids from Lgr5^+^ intestinal stem cells was a breakthrough in tissue culture technology (4). Since then, three-dimensional (3D) intestinal enteroids have been widely used as models to study human gut physiology and pathophysiology as well as host-microbe interactions (5–7). Not only do enteroids render a truer physiological representation of the human epithelium, but they also offer a practical and reliable system to probe mechanisms and interventions at the gut mucosal interface. The 3D spheroid conformation can be adapted to produce a 2D monolayer configuration with enteroids seeded on a semipermeable membrane (i.e., Transwell insert) (8–10). An important practical advantage to this simplified format is that it allows for direct and controlled access to the apical (mimicking the lumen) and basolateral side of the epithelial cells, thus facilitating experimental manipulation and evaluation of outcomes. Enteroid monolayers, which can be generated from any gut segment, reflect the undifferentiated (crypt-like) and differentiated (villus-like) intestinal epithelial cell composition (i.e., absorptive enterocytes, goblet cells, enteroendocrine cells, and Paneth cells) (7, 11–14) of the human gut and exhibit segment-specific phenotypic (15) and functional attributes of the normal gastrointestinal epithelium including production and secretion of mucus (9, 16). To better recreate the cellular complexity of the gastrointestinal mucosal barrier, we devised a human primary cell co-culture system consisting of enteroid monolayers and macrophages seeded on the basolateral side (11). Studies using this enteroid-macrophage co-culture model demonstrated physical and cytokine/chemokine-mediated interactions between intestinal epithelial cells and macrophages in the presence of pathogenic *Escherichia coli* (11, 17). Aiming to expand this co-culture conformation to include other phagocytic cells, we established an *ex vivo* co-culture model containing intestinal epithelial cells and human primary polymorphonuclear neutrophils (PMN) facing the monolayers’ basal membrane. The histological and functional features of this co-culture model, such as cell integration, PMN phenotype, PMN-epithelial cell physical and molecular interactions, and cell function were characterized. Coordinated epithelial and PMN anti-microbial response was examined using *Shigella* as model enteric pathogen. *Shigella* causes diarrhea and dysentery in humans by trespassing the colonic barrier via M cells and infecting epithelial cells, and this process involves massive recruitment of PMN (18). The human enteroid-PMN co-culture revealed dynamic changes in PMN phenotype triggered by epithelial cell contact in the intestinal environment and modeled a paradoxical role of PMN contributing to inflammation and controlling infection.

## Results

### Establishment of a PMN-enteroid co-culture and PMN-epithelial cell interaction

To interrogate PMN adaptation and function in the human gut, we established an enteroid-PMN co-culture model that contains human, primary, enteroid monolayers and PMN isolated from peripheral blood. A human co-culture containing enteroid monolayers and macrophages with similar configuration has been developed by our group (11). Human ileal 3D organoids derived from Lgr5^+^-containing biopsies from healthy subjects were seeded upon the inner (upper) surface of a Transwell insert and allowed to grow until they reached confluency. PMN isolated from peripheral blood of healthy human adult volunteers and exhibiting a CD15^+^CD16^+^CD14^-^phenotype (19) (Figure 1A) were seeded on the outer (bottom) surface of the insert (Figure 1B). Confocal immunofluorescence microscopy and H&E staining confirmed the co-culture’s expected epithelial cell polarity with the brush border oriented towards the luminal side (apical compartment) and adherent PMN facing the basolateral side of the monolayer (Figure 1B). Interestingly, the basolaterally seeded PMN rapidly mobilized towards the epithelium. Within 30 min, PMN migrated through the insert’s pores and intercalated within the epithelial cells (Figure 1C). The migrating PMN could be retrieved from the apical compartment media; between 1.5 and 2% of the seeded PMN transmigrated across the epithelium (Figure 1D) were calculated. The addition of PMN to the enteroid monolayer modestly increased epithelial permeability (a 21% reduction in transepithelial electrical resistance [TER] was observed), although the difference did not reach statistical significance (Figure 1E). Because IL-8 promotes PMN recruitment (20), we examined the effect of exogenous IL-8 on the mobilization of PMN co-cultured with epithelial cells and on monolayer permeability. Apical treatment of monolayers with 100 ng/ml IL-8 significantly increased PMN epithelial transmigration (1.8-fold) (Figure 1D) and barrier permeability; a 38% decrease in TER as observed in co-cultures containing both PMN and IL-8 (Figure 1F). Importantly, IL-8 alone did not affect the permeability of monolayers (Figure 1F).

**Figure 1.**
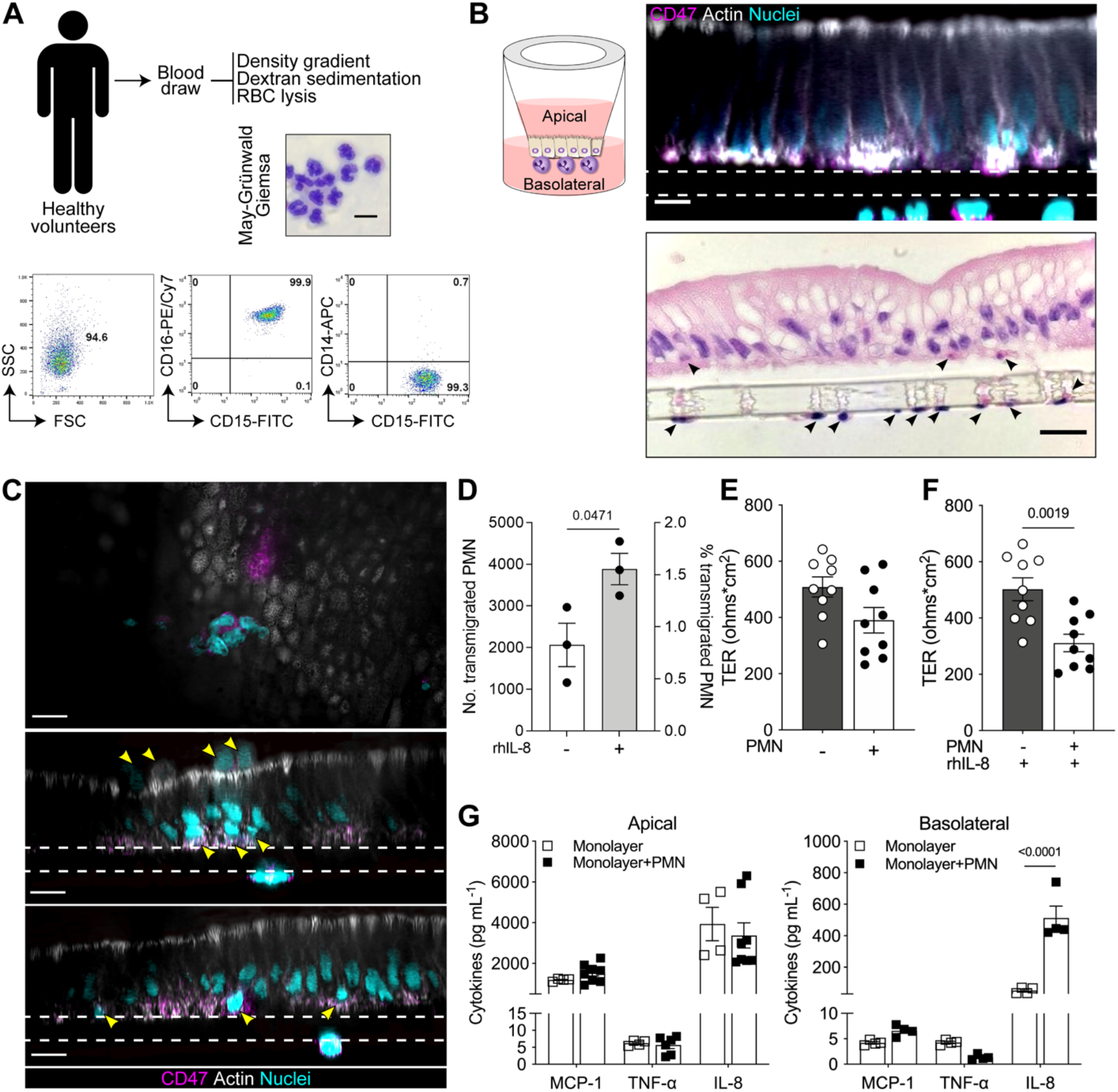
Establishment of a human PMN-enteroid co-culture. (**A**) May-Grünwald Giemsa image of PMN isolated from peripheral blood (top; scale bar = 10 μm). Representative scatter plots of PMN phenotype (bottom). (**B**) Schematic representation the PMN-enteroid co-culture model. Confocal XZ projection (top; Transwell insert, dashed lines; scale bar = 10 μm) and H&E (bottom; scale bar = 20 μm) images of the PMN-enteroid co-culture. Arrowheads indicate PMN seeded on the Transwell insert and at the base of the columnar epithelium. (**C**) Representative immunofluorescence confocal microscopy images (top, XY projection from the apical membrane; middle and bottom, XZ projections) of enteroid monolayers with PMN intercalated and moving up within the epithelial cell monolayer within 30 min of co-culture. Arrowheads indicate PMN. Transwell insert, dashed line. Scale bar = 10 μm. (**D**) Number and proportion of PMN transmigrated to the luminal compartment 2h after being added to the co-culture in the absence or presence of apically delivered rhIL-8. Percent of transmigrated PMN was calculated as the ratio of PMN retrieved in the apical media/number of PMN attached to the Transwell insert (~2.75×10^5^ cells) x100. Each dot represents the average of three replicate wells; data are shown as mean ± S.E.M. from three independent experiments. (**E**) TER of enteroid monolayers (grey bar) and PMN-enteroid co-cultures (white bar) in a 2h co-culture. (**F**) TER of enteroid monolayers and PMN-enteroid co-culture apically treated with rhIL-8 for 2h. (E, F) Each dot represents an independent monolayer; data are shown as mean ± S.E.M. from three independent experiments. (**G**) Cytokines secreted into the apical and basolateral compartments after 2h of co-culture. Data are shown as mean ± S.E.M. from three independent experiments in triplicate. *p* values were calculated by Student’s *t*-test.

Since cell movement is influenced by molecular mediators, we next examined the presence of cytokines (pro- and anti-inflammatory) and chemoattractant molecules in tissue culture media collected from the apical and basolateral compartments of enteroid monolayers alone and of enteroid-PMN co-cultures. Basolateral levels of IL-8 produced by the PMN-containing enteroids were 10-fold higher as compared with IL-8 produced by the monolayers alone, while MCP-1 and TNF -α remained unaffected by addition of PMN (Figure 1G). Similarly, the presence of PMN did not affect apical secretion of MCP-1, TNF-α, and IL-8 (Figure 1G). Production of MCP-1 was distinctly polarized, with higher levels being released to the apical side of the epithelial barrier. IL-1β, IL-6, IL-10, IL-12p70, IFN-γ, and TGF-β1 were measured but determined to be below the limit of detection of the assay for each cytokine.

Taken together, these results demonstrate adequate engraftment of PMN on the basolateral side of the enteroid monolayer, rapid migration of PMN across the monolayer to the luminal side, PMN-induced basolateral secretion of IL-8, and membrane de-stabilization (increased permeability) by IL-8-enhanced PMN transepithelial movement.

### The human intestinal epithelium environment determines PMN immune phenotype and functional capacity

Cell phenotype, morphology, and function can be affected by the surrounding tissue and molecular environment. We hypothesized that the immune phenotype and functional capacity of PMN added to the enteroid monolayer would be influenced by their proximity or direct contact with the intestinal epithelium. To explore this hypothesis, we determined the expression of cell surface markers and phenotypic features of PMN isolated from peripheral blood in comparison with those of PMN within the enteroid co-culture. Two populations of PMN co-cultured with enteroids were investigated: 1. PMN that had been in direct contact with the epithelial cells and traversed across the monolayers, and 2. PMN harvested from the basolateral media as tissue adjacent milieu (Figure 2A). PMN co-cultured with enteroid monolayers had a distinct phenotypic profile as compared with PMN freshly isolated from peripheral blood. Regardless of their location, whether in contact with cells or in basolateral media, PMN co-cultured with enteroid monolayers exhibited increased expression of CD18 (β2 integrin), a molecule that participates in extravasation of circulating PMN, as well as upregulation of CD47, a receptor for membrane integrins involved in cell adhesion and migration, and of CD88 (C5a receptor), a molecule that mediates chemotaxis, granule enzyme release, and super anion production (Figure 2B; Figure S1B). CD66b (CEA cell adhesion molecule 8), a marker of secondary granule exocytosis and increased production of reactive oxygen species, was likewise increased, but only in PMN harvested from the basolateral media (not in contact with cells) (Figure 2C; Figure S1B). In contrast, the expression of CD182 (CXCR2 or IL-8RB) was reduced in all PMN in the co-culture, without distinction of retrieving site (Figure 2C; Figure S1B). PMN that were in close contact with epithelial cells exhibited increased expression of CD15 (E-selectin), a molecule that mediates PMN extravasation; CD16 (FcγRIII), a receptor for IgG that mediates degranulation, phagocytosis, and oxidative burst; and CD11b (α integrin), a protein that facilitates PMN adhesion and, along with CD18, forms the Mac-1 complex implicated in multiple anti-microbial functions (e.g., phagocytosis, cell-mediated cytotoxicity, cellular activation) (Figure 2D; Figure S1B). These results demonstrate that PMN within the intestinal epithelial environment undergo unique phenotypic adaptations, some of which are driven by molecular signals while others require direct PMN-epithelial cell contact.

**Figure 2.**
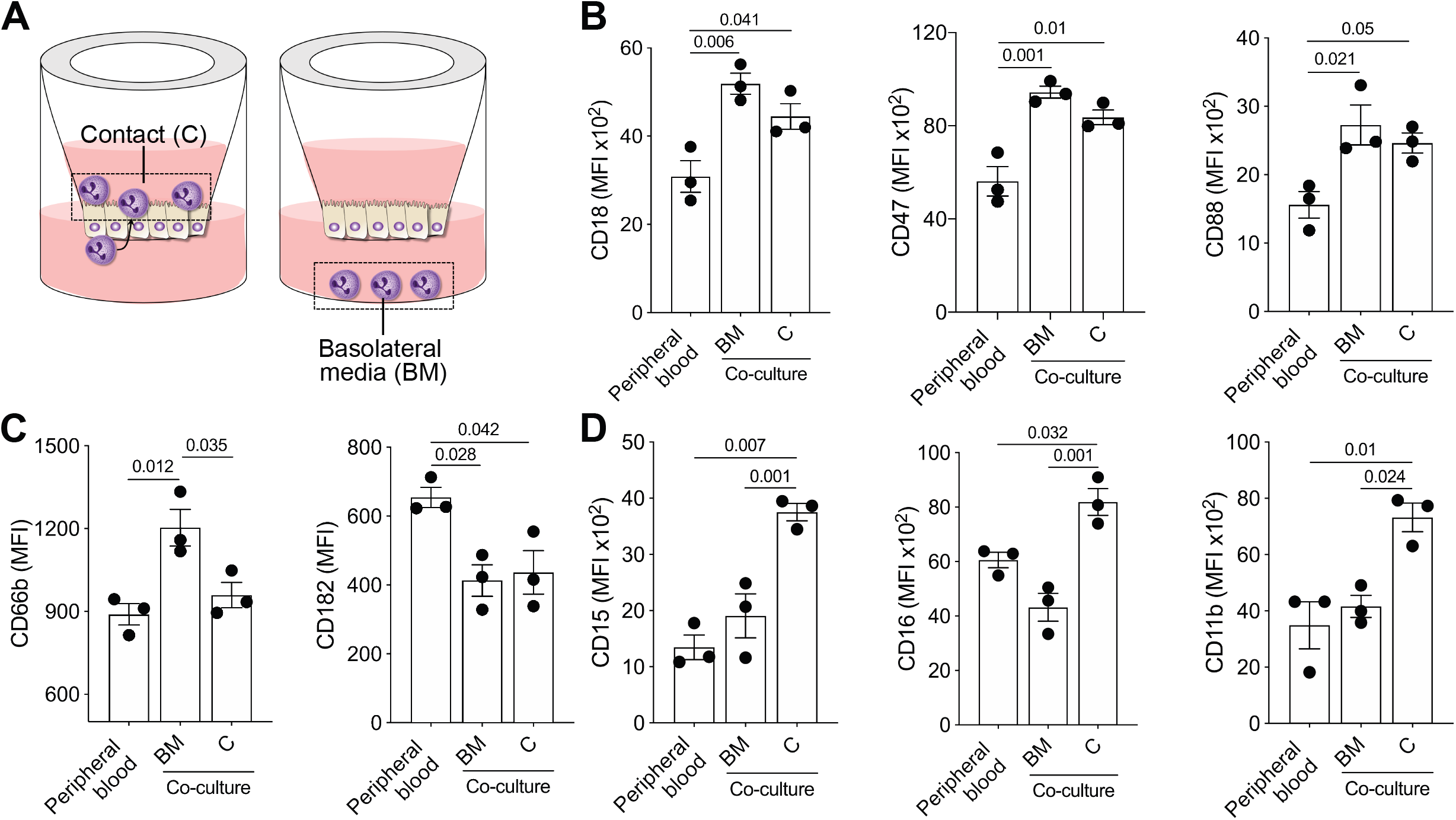
Distinctive phenotype of PMN within the human intestinal environment. (**A**) Schematic representation of PMN in direct contact with the apical membrane of epithelial cells (C) or in the basolateral media (BM). (**B, C, D**) Phenotype of isolated PMN, PMN in C and BM determined by flow cytometry within 2h of co-culture. Each dot represents data collected of three replicate wells; data are shown as mean ± S.E.M. from three independent experiments. *p* values were calculated by one-way-ANOVA with Tukey’s post-test for multiple comparisons. The gating strategy and viability plot of PMN co-cultured with epithelial cells are shown in Figure S1A.

### PMN interaction with *Shigella* as a model enteric pathogen

To interrogate human epithelial cell and PMN interactions in the context of an enteric infection, we exposed the co-culture to wild type (WT) *Shigella flexneri* 2a (strain 2457T) as a model pathogen. PMN participate in *Shigella* pathogenesis through secretion of pro-inflammatory cytokines and deploy anti-microbial functions including phagocytosis, proteolytic enzymes, anti-microbial proteins, and neutrophil extracellular traps (NETs) production. We first determined baseline responses of peripheral blood PMN in the presence of *Shigella*. PMN phagocytosed FITC-labeled *S. flexneri* 2a 2457T within 10 min of exposure (Figure 3A; left panel); bacterial phagocytosis increased over time (up to 1h tested) reaching a maximum effect at 30 min (Figure 3A; middle and right panel). In parallel, the number of bacteria recovered from the culture supernatant of *Shigella*-exposed PMN decreased significantly within 30- and 60-min exposure, in comparison to the number of bacteria recovered from control wells containing *Shigella* alone (in the absence of PMN) resuspended in tissue culture media or media that had been exposed to PMN to control for any soluble bactericidal source (Figure 3B). Overall PMN viability was not affected during the first 2h of *Shigella* exposure but decreased significantly by 3h post infection (Figure 3C; Figure S2). *Shigella-exposed* PMN exhibited changes in cell morphology and motility; formation of pseudopodia (projections of the cell membrane that enable locomotion) was observed within 10 min post infection (Figure 3D). In the presence of *Shigella*, PMN also displayed dynamic amoeboid motility toward the bacteria, and released NETs evidenced by overlap of DNA and histone 3, with entrapped bacilli (Figure 3E). FITC-stained *Shigella* colocalized with PMN phagolysosome (Figure 3F) and the number of *Shigella^+^Lys^+^* PMN increased over time and remained steady up to 2h post infection (Figure 3G). The observation of NET formation at early stages of infection, likely involved a small number of PMN, as the cell population remained largely viable. In addition, the increase in phagocytic activity later during infection is consistent with functional heterogeneity and plasticity of PMN. It has been hypothesized that stepwise deployment of antimicrobial functions by PMN with intrinsically different phenotypes might be beneficial to fine-tune antibacterial responses when faced with an acute infection to prevent collateral damage due to hyperactivation (21).

**Figure 3.**
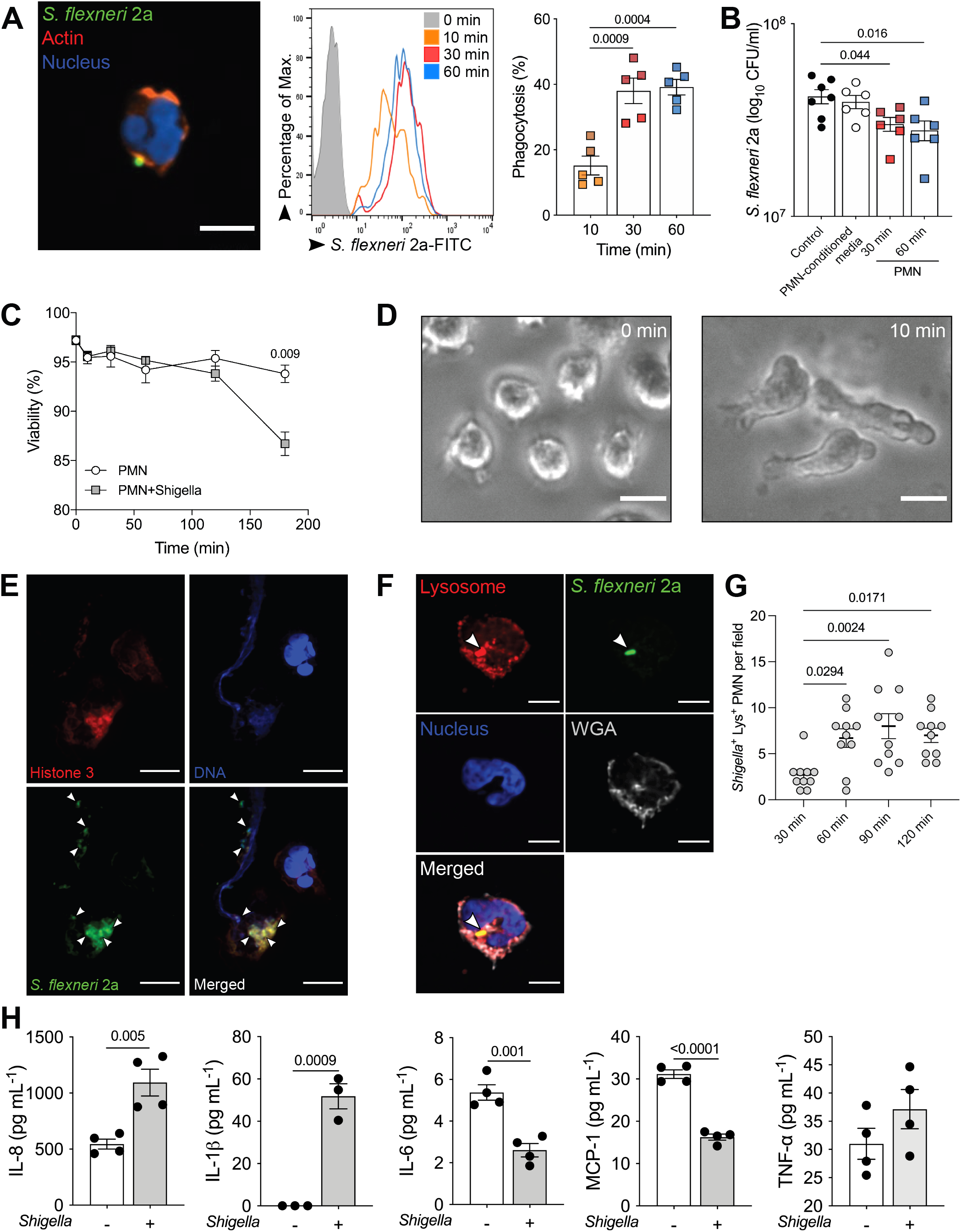
Innate immune responses of human PMN to *S. flexneri*. (**A**) Representative confocal microscopy image (left, XY projection; scale bar = 10 μm), histogram of *S. flexneri* 2a-FITC uptake by human PMN (middle), and percentage of phagocytosis at 10-, 30- and 60-min post infection (left). Each dot represents the average of three replicates and PMN from five individual donors; data are shown as mean ± S.E.M. from three independent experiments. (**B**) Extracellular *S. flexneri* 2a colony forming units (CFU) in culture media following PMN exposure to *S. flexneri* 2a (1×10^8^ CFU/ml) for 30- and 60-min; controls included *S. flexneri* 2a in culture media or in PMN-conditioned media without PMN. Each dot represents the average of three replicates and PMN from six individual donors; data are shown as mean ± S.E.M. from three independent experiments. (A, B) *p* values were calculated by one-way ANOVA with Tukey’s post-test for multiple comparisons. (**C**) PMN viability in the presence and absence of *S. flexneri* 2a. Data represents the average of three replicates and PMN from three individual donors; data are shown as mean ± S.E.M. from three independent experiments. (**D**) PMN morphology before (0 min) and after (10 min) exposure to *S. flexneri* 2a. Scale bar = 10 μm. (**E**) Confocal microscopy image of NETs 30 min after PMN exposure to *S. flexneri* 2a. Arrowheads indicate bacteria. Scale bar = 5 μm. (**F**) Immunofluorescence image of *S. flexneri* 2a-FITC colocalization with PMN lysosome 30 min post infection. Arrowheads indicate bacteria intracellularly and within the lysosome compartment. Scale bar = 5 μm. (**G**) Colocalization of *S. flexneri* 2a-FITC with PMN 30 min to 2h post infection. Each dot represents the number of *Shigella*^+^ Lys^+^ PMN per 10 consecutive microscopy fields. *p* values were calculated by one-way ANOVA with Tukey’s post-test for multiple comparisons. (**H**) Cytokines secreted in culture supernatants of PMN alone and PMN exposed to *S. flexneri* 2a for 2h. Data are shown as mean ± S.E.M. from three independent experiments in triplicate. (C, G) *p* values were calculated by Student’s *t* test.

In addition, PMN upregulated production and secretion of IL-8 and IL-1β, which are key molecular mediators of *Shigella* pathogenesis, 2h post infection, while production of IL-6 and MCP-1 was downregulated; production of TNF-α was not affected (Figure 3H). IL-10, TGF-β1, and IFN-γ were also measured but were below the limit of detection of the assay. These results showed that PMN anti-microbial responses against *Shigella* involved morphological changes, phagocytic activity, and modulation of inflammatory cytokines.

### Coordinated epithelial cell and PMN responses to *Shigella* infection in the co-culture model

We interrogated PMN and epithelial cell interactions and coordinated responses to *Shigella* in the PMN-enteroid co-cultures. PMN were added to the enteroid monolayer as described above (Figure 1B), allowed to settle for 2h, and then apically exposed to WT *S. flexneri* 2a (MOI=10) for another 2h. Non-exposed co-cultures served as controls. Consistent with our previous observation, PMN facing the basolateral side of the epithelial cells moved swiftly through the filter pores, traversed between the epithelial cells and across the monolayer, and protruded on the apical side. PMN basolateral-apical transmigration (both total number and the proportion) increased in the presence of *Shigella* (Figure 4A). Confocal immunofluorescent images revealed PMN phagocytosis of bacteria (Figure 4B; CD47^+^ PMN stained in red, engulfed *S. flexneri* 2a in green, actin in white, and nuclei in blue) and NET formation with trapped organisms (Figure 4C and movie S1) on the luminal surface.

**Figure 4.**
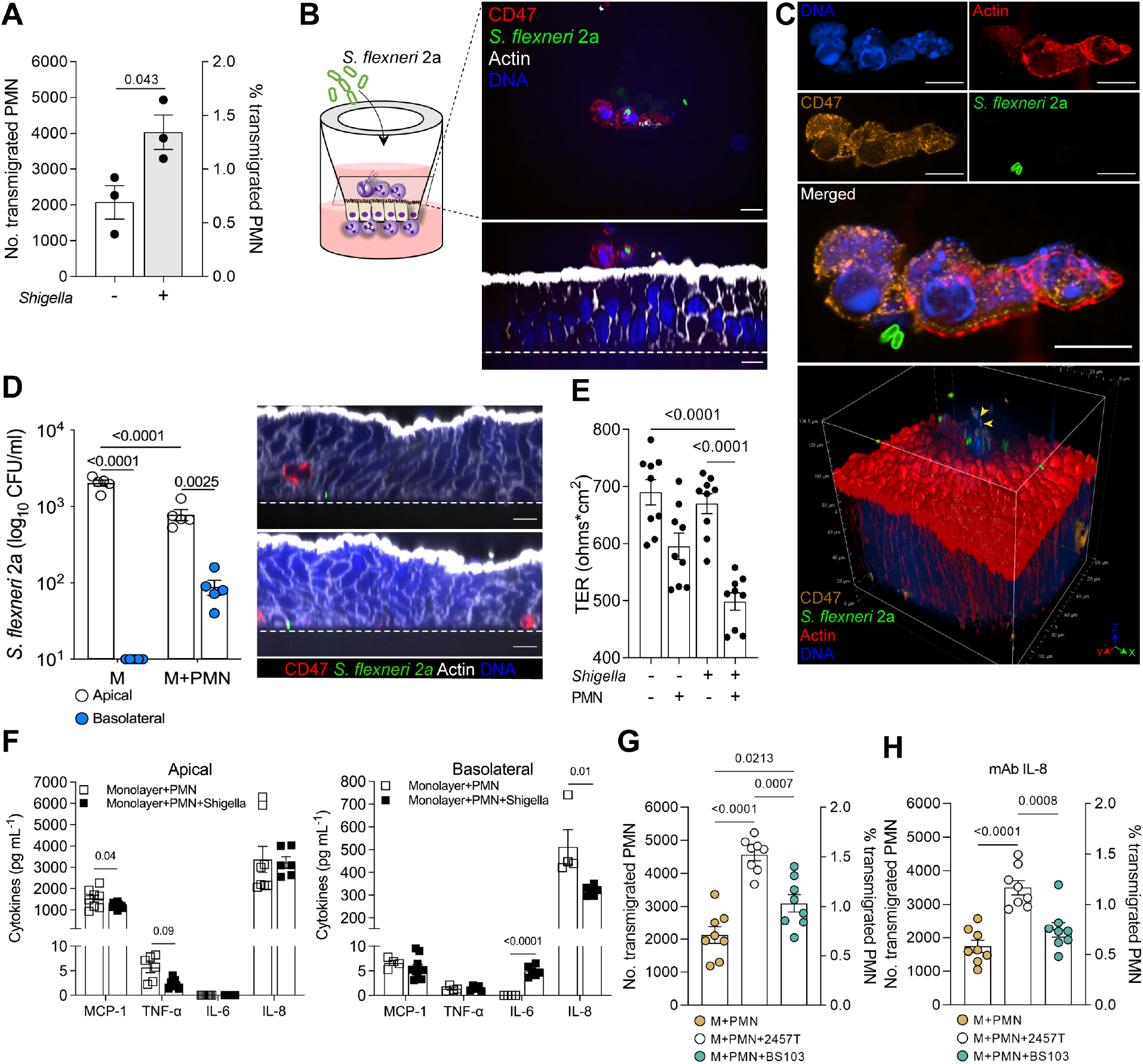
Coordinated innate immune response to *Shigella* by human intestinal epithelial cells and PMN. (**A**) Numbers and proportion of PMN transmigrated to the apical compartment of enteroid monolayers exposed or not to *S. flexneri* 2a WT for 2h. Percent of transmigrated PMN was calculated as the ratio of PMN retrieved in the apical media/number of PMN attached to the Transwell insert (~2.75×10^5^ cells) x100. (**B**) Schematic representation (left) and confocal microscopy images (top, XY; bottom, XZ projections) of PMN-enteroid co-culture infected with *S. flexneri* 2a for 2h. Transwell insert, dashed line. Scale bar = 10 μm. (**C**) Confocal microscopy images and 3D projection of NET on the apical membrane of PMN-enteroid co-culture exposed to *S. flexneri* 2a for 2h. Arrowheads indicate decondensed extracellular thread-like DNA. Scale bar = 10 μm. (**D**) *Shigella* 2a CFU in the apical and basolateral media of enteroid monolayers (M) and PMN-enteroid co-cultures (M+PMN) infected apically for 2h. Confocal microscopy images (XZ projections) of of PMN-enteroid co-culture infected with *S. flexneri* 2a for 2h. Transwell insert, dashed line. Scale bar = 10 μm. (A, D) Each dot represents the mean of three replicate wells; data are shown as mean ± S.E.M. from three (A) and five (D) independent experiments. *p* values were calculated by Student’s *t*-test (A), Student’s *t*-test multiple comparisons (D). (**E**) TER of enteroid and PMN-enteroid co-cultures exposed to *S. flexneri* 2a for 2h; enteroid monolayer alone or co-cultured with PMN for 2h were included as controls. Each dot represents an independent monolayer; data are shown as mean ± S.E.M. from three independent experiments. *p* values were calculated by one-way ANOVA with Tukey’s post-test for multiple comparisons. (**F**) Cytokines secreted in the apical and basolateral media of PMN-enteroid co-cultures exposed or not to *S. flexneri* 2a for 2h. Data are shown as mean ± S.E.M. from three independent experiments in triplicate. *p* values were calculated by Student’s *t*-test. (**G**) Numbers and proportion of PMN transmigrated to the apical compartment of enteroid monolayers exposed or not to *S. flexneri* 2a 2457T or BS103 for 2h. (**H**) Numbers and proportion of PMN transmigrated to the apical compartment of enteroid monolayers pre-treated with human anti-IL-8 monoclonal antibody and exposed or not to *S. flexneri* 2a 2457T or BS103 for 2h. Each dot represents an independent monolayer; data are shown as mean ± S.E.M. from two independent experiments. *p* values were calculated by one-way ANOVA with Tukey’s post-test for multiple comparisons.

The addition of PMN to the enteroid monolayers enabled *Shigella* penetration and cell invasion through the basolateral side (Figure 4D), whereas in the absence of PMN, *Shigella* was unable to trespass the intact enteroid; no bacteria could be recovered from apically exposed enterocytes (Figure 4D). TER values were significantly reduced in the *Shigella-exposed* PMN-enteroid co-culture as compared with *Shigella-exposed* monolayers without PMN or monolayers control (no PMN, no *Shigella)* (Figure 4E). This observation is consistent with tissue damage and loss of barrier function that enables bacterial translocation. Similar to the results shown in Figure 1E, TER values in monolayers containing PMN were somewhat lower compared to monolayers alone, although the difference did not reach statistical significance.

Cytokines produced by the *Shigella*-infected and non-infected PMN-enteroid co-cultures were also examined in the culture media collected from the apical and basolateral compartments. *Shigella* infection reduced apical production of MCP-1 and TNF-α by the co-cultured cells, increased production of IL-6, and substantially reduced secretion of IL-8 basolaterally (Figure 4F). IL-6 was only detected in *Shigella-exposed* PMN-enteroid co-cultures and secreted exclusively to the basolateral side (Figure 4F). IL-1β, a hallmark of *Shigella* pathogenesis, was measured but found to be below detectable levels in both apical and basolateral compartments.

Next, we examined whether PMN migration across the epithelial monolayer was dependent on bacterial virulence factors. To this end, PMN-enteroid co-cultures were exposed to *S. flexneri* 2a WT strain 2457T (used in previous experiments) or *S. flexneri* 2a BS103, a virulence plasmid-curated strain (22). The number and proportion of PMN retrieved from the apical compartment increased significantly in co-cultures infected with either strain as compared with non-infected controls (Figure 4G; Figure S3A). However, PMN transmigration was higher in response to the virulent as compared to the non-invasive strain (Figure 4G).

To evaluate whether PMN transmigration upon infection was influenced by a molecular (cytokine) gradient, PMN-enteroid co-cultures were pre-treated with anti-IL-8 monoclonal antibody and infected with *S. flexneri* 2a 2457T or BS103 as described above. Blocking of IL-8 in the apical and basolateral culture media was confirmed by ELISA (Figure S3B). PMN basolateral-to-apical transmigration still increased in cultures infected with *Shigella* WT in IL-8 blocking conditions (although the proportion of transmigrated PMN was lower), but not in those exposed to the plasmid-curated *Shigella* BS103 strain (Figure 4H).

Collectively, these observations demonstrate that *Shigella* infection causes active recruitment of PMN from the basolateral side of the epithelium (where they were seeded) to the luminal side, and this process involves bacterial virulence factors and host molecular mediators such as epithelial-cell derived IL-8. Paradoxically, the PMN activation and transmigration produced epithelial barrier damage that enabled *Shigella* penetration and basolateral infection; meanwhile PMN actively engulfed bacteria and increased production and secretion of inflammatory cytokines to the apical and basolateral compartments.

### Phenotypic changes in PMN co-cultured with epithelial cells in response to *Shigella* infection

Additional experiments were conducted to determine the phenotypic features of PMN co-cultured with epithelial cells upon *Shigella* infection. PMN co-cultured with intestinal epithelial cells and exposed to *Shigella* exhibited increased expression of CD88, CD47, and CD66b as compared with PMN in co-cultures that remained uninfected (Figure 5A; Figure S4B). In contrast, CD15 and CD18 expression was decreased on PMN from *Shigella*-infected co-cultures (Figure 5B; Figure S4B). CD16, CD11b, and CD182 remained unchanged (Figure 5C; Figure S4B). These results suggest that PMN residing within the intestinal environment undergo immune phenotypic adaptation as a result of pathogen exposure consistent with increased anti-microbial function (i.e., activation and expression of molecules that facilitate chemotaxis and epithelial transmigration).

**Figure 5.**
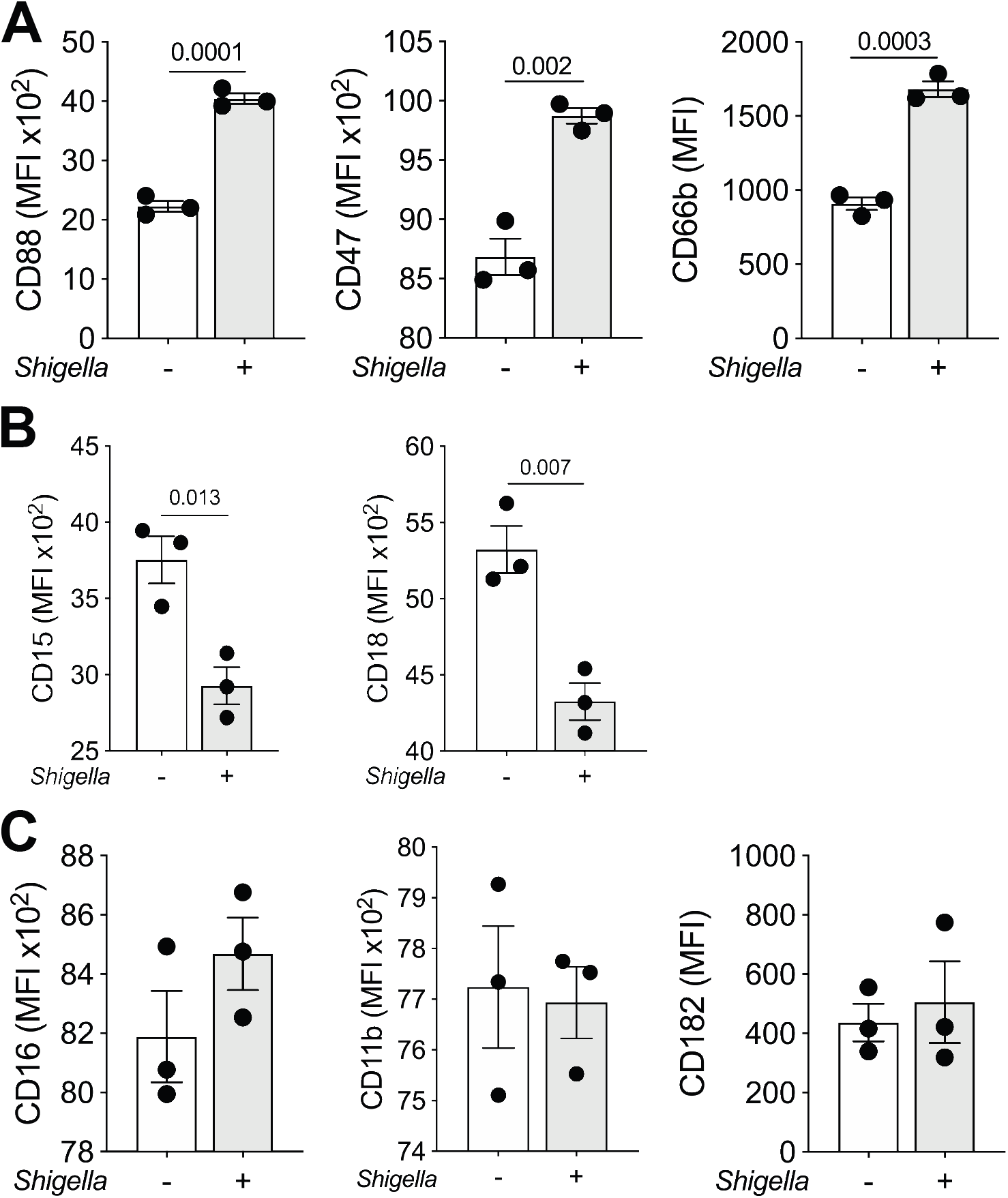
Immune phenotype of PMN co-cultured with epithelial cells and exposed to *Shigella*. (**A, B, C**) Cell surface expression of CD15, CD16, CD11b, CD18, CD88, CD66b, CD47, and CD182 on PMN embedded in the enteroid monolayer (co-culture) exposed or not to *S. flexneri* 2a for 2h. Each dot represents data collected of three replicate wells; data are shown as mean ± S.E.M. from three independent experiments. *p* values were calculated by Student’s *t*-test. The gating strategy and viability plot of PMN co-cultured with epithelial cells and exposed to *S. flexneri* 2a WT are provided in Figures S4.

## Discussion

Epithelial cells and innate phagocytic cells underlying the intestinal epithelium work synergistically, preventing the trespassing of harmful agents and deploying rapid and effective host defense mechanisms against pathogens. PMN are the first innate immune cells recruited in response to gastrointestinal tissue inflammation and infection and they play a critical role in initiating host immune responses (23). Patients with neutrophil disorders are prone to recurrent microbial infection (24, 25). In this manuscript, we report the successful establishment of an *ex vivo* primary human intestinal epithelial-PMN co-culture, and we describe cell interactions, phenotypic and functional adaptations, and cellular and molecular innate responses to *Shigella* as a relevant dysenteric pathogen.

Our group developed the first *ex vivo* human enteroid and monocyte-derived macrophage co-culture model in a monolayer format (11). The same approach was used to produce the PMN-enteroid co-culture described herein. Differently from macrophages, which remained in the basolateral side of the monolayer (where seeded) and responded to luminal organisms by extending transepithelial projections between adjacent epithelial cells, PMN rapidly migrated from the basolateral side of the epithelial cell and across the monolayer via the paracellular space. Histological and confocal microscopy images revealed PMN crawling through the Transwell insert pores, embedding at the base of the epithelial cells, and emerging on the luminal side of the enteroid monolayer, all within a few hours of co-culture. While macrophages contributed to cell differentiation and stabilized the epithelial barrier in our previous studies, PMN transmigration resulted in increased barrier permeability that enabled bacteria invasion.

The capacity of PMN to migrate across the vascular endothelium (26) and a variety of tissues (27, 28), including epithelial cell layers (29–32), has been documented *in vivo* (mainly in animal models) or *in vitro* using cell lines. These processes have been associated with PMN activation as a result of microbial sensing, inflammation, or danger signals. The level of myeloperoxidase (MPO), one of the principal enzymes contained in PMN granules and released upon activation is, in fact, in stool, a biomarker of inflammatory bowel disease severity (33). In our human PMN and intestinal epithelial cell co-culture model, PMN migrated even in the absence of external stimulatory signals. PMN are not typically present in the homeostatic gut but actively recruited by signs of inflammation or infection (34); therefore, unprovoked PMN migration would not be expected.

The tissue microenvironment can influence immune cell phenotype and effector function capabilities (35, 36). PMN from peripheral blood exhibited rapid phenotypic changes when incubated with enteroid monolayers. They acquired an activated phenotype that was triggered by tissue-derived signals. Compared to the PMN from peripheral blood, PMN co-cultured with epithelial cells had increased expression of CD18, which favors cell binding; of CD88, which is the receptor for C5a and facilitates degranulation and chemotaxis; and of CD47, a cell surface glycoprotein that supports transmigration across endothelial and epithelial cells (37–39). PMN phenotypic changes were influenced by their spatial location, i.e., whether they were in direct contact with epithelial cells or simply present in basolateral culture media. PMN in close proximity with the epithelium (the migratory PMN) had upregulated expression of CD15, which participates in chemotaxis and extravasation from circulation; as well as FcγRIII (CD16), a low affinity Fc receptor for IgG; and CD11b, a marker of cell adhesion and anti-microbial function (phagocytosis, degranulation, oxidative burst) (40, 41). On the other hand, PMN retrieved from the basolateral media, and which had not been in contact with the epithelial layer, exhibited increased expression of CD66b, indicative of PMN activation and degranulation (42). This is to our knowledge the first detailed description of dynamic changes of PMN immune phenotype in a translationally relevant model of the human intestinal epithelium.

MCP-1/CCL2, a chemoattractant and enhancer of bacterial killing and survival of phagocytic cells and IL-8, a hallmark product of intestinal epithelial cells and a potent activator and chemoattractant for PMN (43, 44), were abundantly produced by the ileal monolayers in our co-culture model. TNF-α, a recruiter and activator of phagocytic cells (45) was also detected, albeit at lower levels. IL-8 was further sourced by the PMN within the co-culture and released to the basolateral media; *in vivo* this subepithelial surge of IL-8 likely contributes to PMN recruitment *in vivo*. Intriguingly, expression of IL-8 receptor (CD182/CXCR2) was downregulated in PMN co-cultured with enteroid monolayers as compared to PMN from peripheral blood, most likely reflecting a compensatory mechanism. Still, IL-8 may act via another high affinity receptor (i.e., CXCR1 or IL-8RA) (46, 47). Expression of MCP-1, IL-8, and TNF-α has been reported in intestinal tissue of healthy adults in steady state (48, 49).

Again, clear differences in innate immune functions emerged when comparing cytokine profiles in PMN- and macrophage-enteroid co-cultures. Macrophages contributed to high levels of IFN-γ and IL-6 (11); however, these cytokines were not detected in co-cultures containing PMN. Accordingly, both models were capable of discerning distinct morphological as well as functional phagocytic cell adaptation.

*Shigella* invades the human colon and rectal mucosa and causes severe inflammation, massive recruitment of PMN, and tissue destruction (18). Bloody/mucous diarrhea (dysentery) with large numbers of PMN in stool are hallmarks of shigellosis (50). Human intestinal enteroids can be infected with *Shigella* (51, 52). Hence, our model was fitting to interrogate coordinated innate responses of epithelial cells and PMN to this enteric pathogen. *Shigella* added to the apical side of the enteroid monolayers increased basolateral-to-apical PMN migration. Early efflux of PMN into the colonic tissue has been observed during shigellosis in the infected rabbit loop model (53). Sansonetti and colleagues reported *Shigella*-induced PMN transmigration that promoted invasion of colonic T84 cell (54). The same group later showed that the *Shigella* lipopolysaccharide transcytosed to the basal side of T84 cells enhanced adherence of subepithelial PMN through IL-8 signaling (55). In this manuscript we demonstrated that IL-8 enhanced PMN transepithelial migration across human intestinal epithelial cells in culture. However, the fact that PMN still migrated under IL-8 blocking conditions indicates that signals other than IL-8 are likewise involved in this process. In a series of articles, McCormick and colleagues interrogated molecules and bacterial components involved in *Shigella*-PMN responses in T84 cells (56–59); the group reported the contribution of hepoxilin A3, a product of the cleavage of arachidonic acid via 12/15-lipoxygenase (56) and the requirement of *S. flexneri* plasmid-encoded virulence effectors for PMN migration (60–62). In contrast, Sansonetti et al. found intermediate PMN migration in T84 monolayers exposed to a non-invasive (plasmid curated) *Shigella* strain (54). Our results align with the latter and confirm that virulence factors contribute but are not the only determinants of PMN transepithelial migratory signaling.

During *Shigella* dysentery, PMN migration through the colonic epithelium destabilizes the epithelial barrier and allows massive entry of bacteria into the submucosa with further amplification of infection and tissue destruction (63). This process was recreated in our co-culture model; PMN transmigration across the intestinal monolayer disrupted the polarized epithelial barrier and enabled bacterial invasion. Focal breakdown of the epithelial cell surface has been attributed to PMN migration in various disease states including infectious enterocolitis (64). Intestinal epithelial repair events (e.g., cell proliferation and migration, and closure of leaking epithelial lateral spaces) reportedly begin minutes after acute mucosal barrier injury (65). As a corollary of these observations, we are investigating epithelial repair subsequent to PMN-induced inflammation and cell disruption, and the mechanisms and elements involved.

Modeling the human PMN-epithelial cell interaction *ex vivo*, our findings challenge the notion that M cells are necessary for *Shigella* epithelial translocation and invasion, and hint that the infiltration and barrier disruption by PMN offer an alternative mechanism by which *Shigella* and other invasive enteric pathogens access the host internal compartment. We are presently studying reverse transmigration of bacteria-loaded PMN (out of the lumen and back to the basolateral side) as a possible means to initiate adaptive immunity through cross-presentation. The PMN-enteroid co-culture model is also useful to interrogate molecules involved in PMN activity in the context of enteric infections and to identify targets to prevent intestinal injury and inflammation.

Although acting in a “brute force” manner, PMN deployed potent anti-microbial activity against *Shigella*. As expected, circulating PMN entrapped bacteria into NET structures and exhibited *Shigella* phagocytic activity. PMN anti-microbial functions coincided with increased production of IL-8 and the pyroptosis-inducer IL-1β, and downregulation of IL-6 and MCP-1. Likewise, PMN embedded within the epithelial cells promptly mobilized upon sensing *Shigella* on the epithelial cell surface; PMN that had traversed to the apical side formed NET structures that trapped bacteria and exhibited robust phagocytic and killing capacity. Intriguingly, IL-8 levels were reduced, IL-1β was absent, and IL-6 was increased in the infected PMN-epithelial co-culture as compared to non-infected. A reduction of IL-8 production had been reported in *Shigella*-infected human colonic explants, which was ascribed to anti-inflammatory bacterial proteins or death of IL-8 secreting cells (66). Reduced levels of these inflammatory cytokines may also reflect a negative feedback to prevent further tissue damage. Heightened levels of IL-6 and reduced TNF-α during infection may suggest a protective epithelial mechanism after injury (67, 68). In addition, IL-6 has been attributed a beneficial role in enhancing Th17-protective immunity against *Shigella* re-infection (69). Because cytokines measured in the co-culture supernatant represent the total amount produced by diverse cell types, this readout is limited in its capacity to discern subtle differences between culture conditions.

The immune phenotype of PMN in the co-culture adapted again as a result of *Shigella* infection, with further increases in activation/granule-associated markers CD66b, CD88, and CD47. CD47 has been implicated in PMN paracellular migration through epithelial cells in response to bacterium-derived leukocyte chemoattractant N-formyl-methionyl-leucyl-phenylalanine, in a process that involves intracellular distribution and increased CD47 cell surface expression (38). CD47-deficient mice have increased susceptibility to *E. coli* as a result of reduced PMN trafficking and bacterial killing activity (70). This finding is consistent with our observed upregulation of CD47 in *Shigella-exposed* PMN, which is likely associated with PMN’s anti-microbial activity. CD16 and CD11b expression were unaltered on PMN co-cultured with intestinal enteroids and exposed to *Shigella*, indicating a preserved phagocytic capacity, whereas extravasation and cell adhesion markers CD15 and CD18 were downregulated. It has been reported that CD47 expression increases gradually and modulates CD11b-integrin function and CD11b/CD18 surface expression on PMN, suggesting a regulatory mechanism between these molecules (38, 71). Expression of CD47 is self-protective; it avoids clearance by phagocytic cells (72). The exact role of CD47 expression on PMN during *Shigella* infection remains to be elucidated.

Human intestinal xenografts in immunodeficient mice have been used to model interactions of *Shigella* with the human intestine *in vivo*. The model failed to discern any role of PMN in ameliorating or exacerbating disease but revealed larger number of intracellular bacteria in PMN-depleted mice; the authors concluded that while PMN may contribute to tissue damage, they are important in controlling bacteria dissemination. The combination of species, immunodeficient background, and impracticality are major confounders/limitations of this model (73).

Our study contributed new insights into the morphological, phenotypic and functional adaptation of PMN in the gastrointestinal environment, the close communication between PMN and epithelial cells and their coordinated responses to *Shigella* as a model enteric pathogen. The human PMN-enteroid co-culture described here provides a translationally relevant *ex vivo* model to study human epithelial cell-PMN physiology and pathophysiology, as well as host cell interactions and innate responses to enteric organisms. This model could be useful to interrogate innate immune defense mechanisms to enteric pathogens and to support the development and evaluation of preventive or therapeutic tools.

## Materials and methods

### Human PMN isolation

Human peripheral blood was collected in EDTA tubes (BD Vacutainer) from healthy volunteers enrolled in University of Maryland Institutional Review Board (IRB) approved protocol #HP-40025-CVD4000, and methods were conducted in compliance with approved Environmental Health and Safety guidelines (IBC #00003017). PMN were obtained by Ficoll-Paque (PREMIUM solution, GE Healthcare Bio-Sciences AB, Sweden) gradient centrifugation following dextran (Alfa Aesar, USA) sedimentation (74). Contaminating erythrocytes were removed by hypotonic lysis. After washing, cells were suspended in enteroid differentiation media (DFM) without antibiotics and immediately used. The cell suspension contained >95% of PMN as determined by flow cytometry and May-Grünwald-Giemsa stained cytopreps. PMN viability was >98%. Cell counts were determined using the Guava ViaCount Reagent (Luminex, USA); viable and non-viable cells were distinguished based on differential permeabilities of two DNA-binding dye. Cells were stained following the manufacturer’s instruction and analyzed in Guava 8HT using Viacount software (Luminex, USA).

### Preparation of enteroid monolayers

Human enteroid cultures were established from biopsy tissue obtained after endoscopic or surgical resection from healthy subjects at Johns Hopkins University under Johns Hopkins University IRB approved protocol #NA-00038329, as previously described (14). Briefly, enteroids generated from isolated intestinal crypts from ileal segments were maintained as 3D cysts embedded in Matrigel (Corning, USA) in 24-well plates and cultured in Wnt3A-containing non-differentiated media (NDM) (74). Multiple enteroids were harvested with Culturex Organoid Harvesting Solution (Trevigen, USA), and small enteroid fragments were obtained to create 2D monolayers by digestion with TrypLE Express (ThermoFisher) in 37°C water bath for 90 seconds. Enteroid fragments were resuspended in NDM containing 10 μM Y-27632 and 10 μM CHIR 99021 inhibitors (Tocris) (NDM+inhibitors). The inner surface of Transwell inserts (3.0 μm pore transparent polyester membrane) pre-coated with 100 μl of human collagen IV solution (34 μg/ml; Sigma-Aldrich, USA) were seeded with 100 μl of an enteroid fragment suspension, and 600 μl of NDM+inhibitors was added to the wells of a 24-well tissue culture plate and incubated at 37°C, 5% CO2, as previously described (74). NDM without inhibitors was replaced after 48h, and fresh NDM was added every other day; under these conditions, enteroid cultures reached confluency in 14-16 days. Monolayer differentiation was induced by incubation in Wnt3A-free and Rspo-1-free DFM without antibiotics for 5 days (11). Monolayer confluency was monitored by measuring TER values with an epithelial voltohmmeter (EVOM^2^; World Precision Instruments, USA). The unit area resistances (ohm*cm^2^) were calculated according to the growth surface area of the inserts (0.33 cm^2^).

### PMN-enteroid co-culture

Differentiated enteroid monolayers seeded on Transwell inserts were inverted and placed into an empty 12-well plate. PMN (5×10^5^ in 50 μl of DFM) were added onto the bottom surface of the inserts, and cells were allowed to attach for 2h at 37°C, 5% CO_2_ (inserts remained wet throughout this process). The inserts were then turned back to their original position into a 24-well plate, and DFM was added to the insert (100 μl) and into the well (600 μl). Approximately 45% of the added PMN remained attached to the Transwell insert. TER measurements were collected after 2h, allowing monolayer recovering. For the IL-8 neutralization experiments: 0.4 μg/ml of human IL-8 monoclonal antibody (R&D Systems, USA) was added to the apical and basolateral compartment of enteroid monolayers and to the PMN suspensions before they were co-cultured. This dose of anti-IL8 was approximately 100-fold above the highest value of IL-8 detected in the tissue culture media of *Shigella-exposed* PMN-enteroid co-cultures. The absence of IL-8 in enteroid and PMN culture media was confirmed by electrochemiluminescence ELISA as described in the cytokine measurement section below.

### *Shigella flexneri* 2a strains and infection

*Shigella flexneri* 2a WT strain 2457T and avirulent plasmid-curated strain BS103 were grown from frozen stocks (−80°C) on Tryptic Soy Agar (TSA) (Difco BD, USA) supplemented with 0.01% Congo Red dye (Sigma-Aldrich) overnight at 37°C. Bacterial inoculum was made by resuspending single red (2457T) and white (BS103) colonies in sterile 1X PBS pH 7.4 (Quality Biological), respectively. Bacterial suspension was adjusted to the desired concentration (~1×10^8^ CFU/ml) in advanced DMEM/F12 without serum. A bacterial suspension containing ~5×10^6^ CFU in 50 μl was added directly to PMN (for 30-60 min) or to the apical compartment of enteroid monolayers alone or PMN-enteroid co-culture (for 2h), a multiplicity of infection of 10 relative to 1 PMN.

### PMN transmigration, apical and basolateral harvesting

Basolateral-to-apical PMN transmigration was quantified by measurement of PMN MPO using a commercial kit (Cayman Chemical, Ann Arbor, MI) as previously described (75). The assay was standardized with a known number of human PMN. MPO activity in lysates of enteroid monolayer alone was negligible. Transmigrated PMN were also confirmed by counting cells on the apical membrane of enteroid monolayers of 25 consecutive immunofluorescence microscopy fields. For IL-8 gradient experiments, 100ng/ml of recombinant human (rh) IL-8 (Biolegend, San Diego, CA) were added to the apical side of the enteroid monolayers (76), and PMN transmigration determined as described above.

PMN that migrated across the monolayer were collected from the apical media by washing three times the apical compartment with 1X PBS at room temperature (RT). PMN that were not in contact with the epithelial cells were harvested from the basolateral tissue culture media.

### PMN phagocytosis

*S. flexneri* 2a WT cultures grown overnight as described above were washed, resuspended in sterile PBS and incubated with FITC (Sigma-Aldrich) (20 μg/ml) for 30 min at 37°C. The bacteria suspension was thoroughly washed and adjusted to ~10^8^ CFU/ml in sterile PBS/glycerol (1:2) and stored at −80°C until used. The day of the assay, FITC-labeled *Shigella* was incubated with PMN-autologous human sera for 30 min at 37°C. Opsonized bacteria 5×10^6^ CFU was added to PMN suspension (5×10^5^ cells) and incubated for 10-, 30-, and 60-min. Phagocytosis was measured by flow cytometry. External fluorescence was blocked with the addition of trypan blue, and the difference between MFI blocked and non-blocked samples was used to calculate % phagocytosis (77, 78).

### H&E and immunofluorescence staining

PMN-enteroid co-culture cells were fixed in aqueous 4% paraformaldehyde (PFA; Electron Microscopy Sciences, USA) at RT for 45 min and then washed with PBS. For H&E staining, monolayers were kept for at least 48h in formaldehyde solution, then embedded in paraffin, sectioned, mounted on slides, and stained with H&E. For immunofluorescence, cells were permeabilized and blocked for 30 min at RT in PBS containing 15% FBS, 2% BSA, and 0.1% saponin (all Sigma-Aldrich, USA). Cells were rinsed with PBS and incubated overnight at 4°C with primary antibodies: mouse anti-CD16 (LSBio, USA) and rabbit anti-*S. flexneri* 2a (Abcam, USA) diluted 1:100 in PBS containing 15% FBS and 2% BSA. Stained cells were washed with PBS and incubated with secondary antibodies: goat anti-mouse AF488 and goat anti-rabbit AF594 (both Thermo Fisher Scientific, USA) diluted 1:100 in PBS 1h at RT; phalloidin AF594 or AF633 (Molecular Probes, Thermo Fisher Scientific) was included in this step for actin visualization. Cells were washed and mounted in ProLong Gold Antifade Reagent with DAPI (Cell Signaling Technology, USA) for nuclear staining and maintained at 4°C. Lysosome was stained with LysoTracker™ Red DND-99 (Thermo Fischer Scientific) following the manufacturer’s instructions.

### Immunofluorescence microscopy

Confocal imaging was carried out at the Confocal Microscopy Facility of the University of Maryland School of Medicine using a Nikon W1 spinning disk confocal microscope running NIS-Elements imaging software (Nikon). Images were captured with a 40X or 60X oil objective and settings were adjusted to optimize signal. Immunofluorescence imaging (Figure 3F) was carried out using EVOS FL Imaging systems (fluorescent microscope) at 40X objective lens. Images were collated using FIJI/ImageJ (NIH). Signal processing was applied equally across the entire image. Color channels for Figures 1B and 1C, and actin (Figure 4B) were arranged for contrast purpose.

### Flow cytometry

PMN phenotype was determined using the following human specific monoclonal antibodies from BD Pharmingen: HI98 (anti-CD15, FITC-conjugated), M5E2 (anti-CD14, APC-conjugated), 3G8 (anti-CD16, PE/Cy7-conjugated), D53-1473 (anti-CD88, BV421-conjugated), and BioLegend: TS1/18 (anti-CD18, PE/Cy7-conjugated), ICRF44 (anti-CD11b, BV421-conjugated), 5E8/CXCR2 (anti-CD182, APC-conjugated), CC2C6 (anti-CD47, PE/Cy7-conjugated), HA58 (anti-CD54, APC-conjugated), and G10F5 (anti-CD66b, Pacific Blue-conjugated). All the stains were performed in dark. For all experiments, PMN were stained with 2 μl of Zombie Aqua dye (BioLegend) diluted 1:100 for 15 min at RT. PMN were washed and blocked with 2% normal mouse serum (Thermo Fisher Scientific) for 15 min at 4°C. After washing, cells were resuspended in FACS buffer (PBS containing 0.5% BSA and 2 mM EDTA; all Sigma-Aldrich), and 100 μl of equal number of cells were dispensed in several tubes and stained with antibodies for 30 min at 4°C. Antibodies were used diluted 1:2 – 1:1,000; optional amount was determined by in-house titration. Cells were washed in FACS buffer and either analyzed or fixed in 4% PFA for 15 min at 4°C and analyzed the next day. Marker expression was analyzed in a Guava 8HT using Guava ExpressPro software (Luminex, USA) or BD LSRII using FACSDiva software (BD Biosciences, USA) and analyzed with FlowJo software (v10, Tree Star).

### Cytokine and chemokine measurements

Cytokines and chemokines were measured by electrochemiluminescence microarray using commercial assays (Meso Scale Diagnostic, USA) following the manufacturer’s instructions. IFN-γ, IL-1β, IL-6, IL-10, IL-12p70, TNF-α, MCP-1, TGF-β1, and IL-8 were reported as pg mL^-1^contained in the apical and basolateral culture supernatants.

### Statistical analysis

Statistical significances were calculated using the Student’s *t*-test unpaired, one-way or two-way analysis of variance (ANOVA) with Tukey’s post-test as appropriate. Plots and statistical tests were performed using Prism software v9 (GraphPad, San Diego, CA). Treatment comparisons included at least three replicates and three independent experiments. Differences were considered statistically significant at *p*-value ≤ 0.05. Exact *p* values are indicated in each Figure. Results are expressed as mean ± standard error of mean (S.E.M.).

## Acknowledgements

The authors thank Drs. Robert Kaminski for kindly providing *S. flexneri* 2a BS103 (Walter Reed Army Research Institute), and Mark Donowitz for critical reading of the manuscript; and Robin Barnes and Nancy Greenberg, Clinical Research Nurses at University of Maryland, for their assistance in obtaining peripheral blood samples. The authors would like to acknowledge the Integrated Physiology Core of the Hopkins Conte Digestive Disease Basic and Translational Research Core Center (NIH P30 DK-089502 to N.C.Z.) for providing human enteroids and growth factor conditioned media and the University of Maryland School of Medicine Center for Innovative Biomedical Resources, Flow Cytometry Shared Service, and Confocal Microscopy Core (NIH S10 OD026698). This work was supported by NIH/NIAID P01AI125181 Immunology (to M.F.P) and Enteroid (to N.C.Z.) Cores, and 1R01AI117734-01 (to M.F.P).

## Author Contributions

J.M.L-D. designed and performed experiments and conducted data analyses and interpretation. M.D. provided initial assistance with enteroid monolayers. N.C.Z. and M.F.P. participated in experimental design, data analyses and interpretation, and supervised the study. J.M.L-D. and M.F.P. wrote the manuscript. All authors edited the manuscript.

## Supplementary Figures

**Figure S1. Distinctive phenotype of PMN within the human intestinal environment**. (**A**) Representative flow cytometry plots showing PMN viability, and the gating strategy used to determine cell surface expression of CD15, CD16, CD11b, CD18, CD14, CD47, CD54, CD66b, CD88, and CD182 on PMN. (**B**) Representative histograms showing cell surface markers’ expression on PMN isolated from peripheral blood, PMN that had been in direct contact with the enteroid monolayer (C) or PMN that detached and was retrieved in the basolateral media (BM). Data are representative of three independent experiments.

**Figure S2. PMN viability following *Shigella* infection**. Representative flow cytometry plot depicting PMN viability 30- and 180-min post *Shigella* infection; uninfected PMN incubated in the same conditions served as control. Data are representative of three independent experiments.

**Figure S3. Coordinated response to *Shigella* by human intestinal epithelial cells and PMN**. (**A**) Confocal microscopy images (top, XY projections; bottom, XZ projections) of PMN-enteroid co-cultures apically exposed to *S. flexneri* 2a 2457T (left) or BS103 (right) for 2h. Scale bar = 20 μm. (**B**) Total amount of IL-8 in the apical and basolateral media of PMN-enteroid co-cultures pretreated or not with anti-IL-8 monoclonal antibody and exposed or not to *S. flexneri* 2a 2457T or BS103 for 2h. Data are shown as mean ± S.E.M. from two independent experiments in duplicate.

**Figure S4. Immune phenotype of PMN co-cultured with epithelial cell monolayers and exposed to *Shigella*. (A)** Representative flow cytometric PMN live/dead plot and (**B**) contour plots and histograms depicting cell surface markers’ expression on PMN co-cultured with enteroid monolayers and exposed (or not) to *S. flexneri* 2a for 2h; PMN were retrieved from the apical side. Data are representative of three independent experiments.

**Movie S1. NET on the apical cell membrane of PMN-enteroid co-culture exposed to *Shigella*.** Three-dimensional projection of the co-localization of decondensed extracellular thread-like DNA (blue), CD47 (orange) and actin (red) with trapped *S. flexneri* 2a (green).

