## Supplementary figures and images for "Epithelial and neutrophil interactions and coordinated response to *Shigella* in a human intestinal enteroid-neutrophil co-culture model"

### Supplemental Figure 1

#### Slide 1
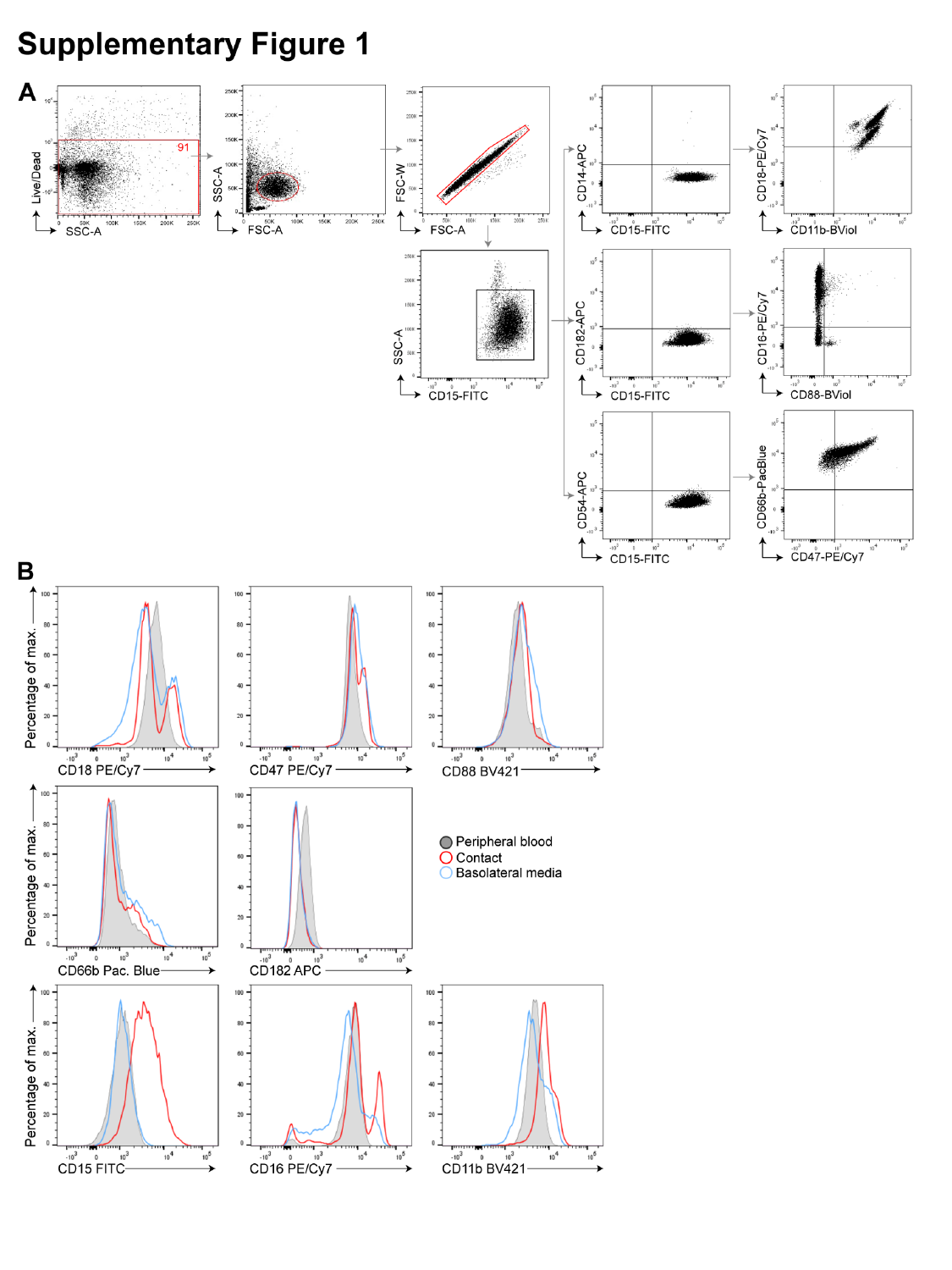

### Supplemental Figure 2

#### Slide 1
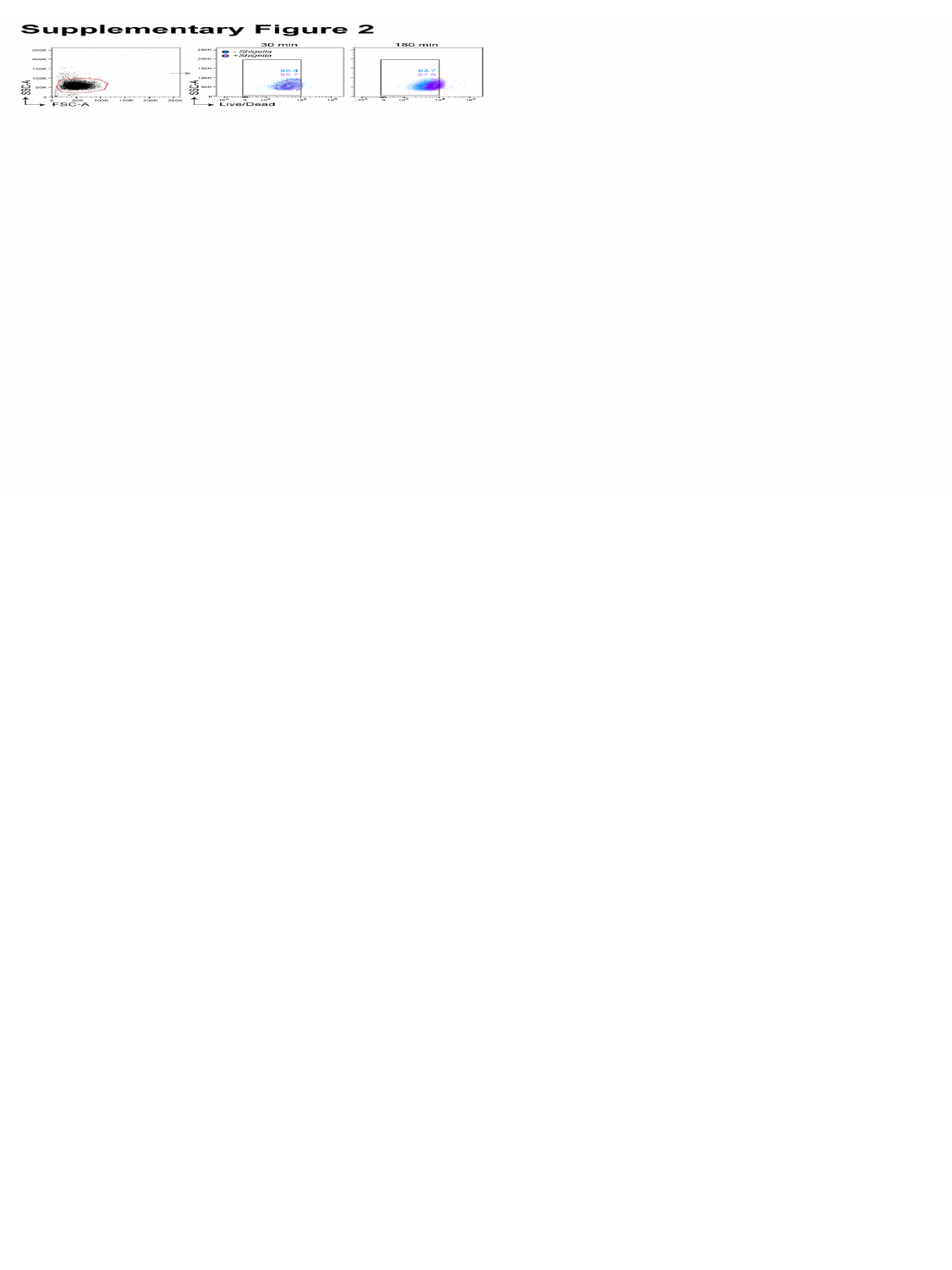

### Supplemental Figure 3

#### Slide 1
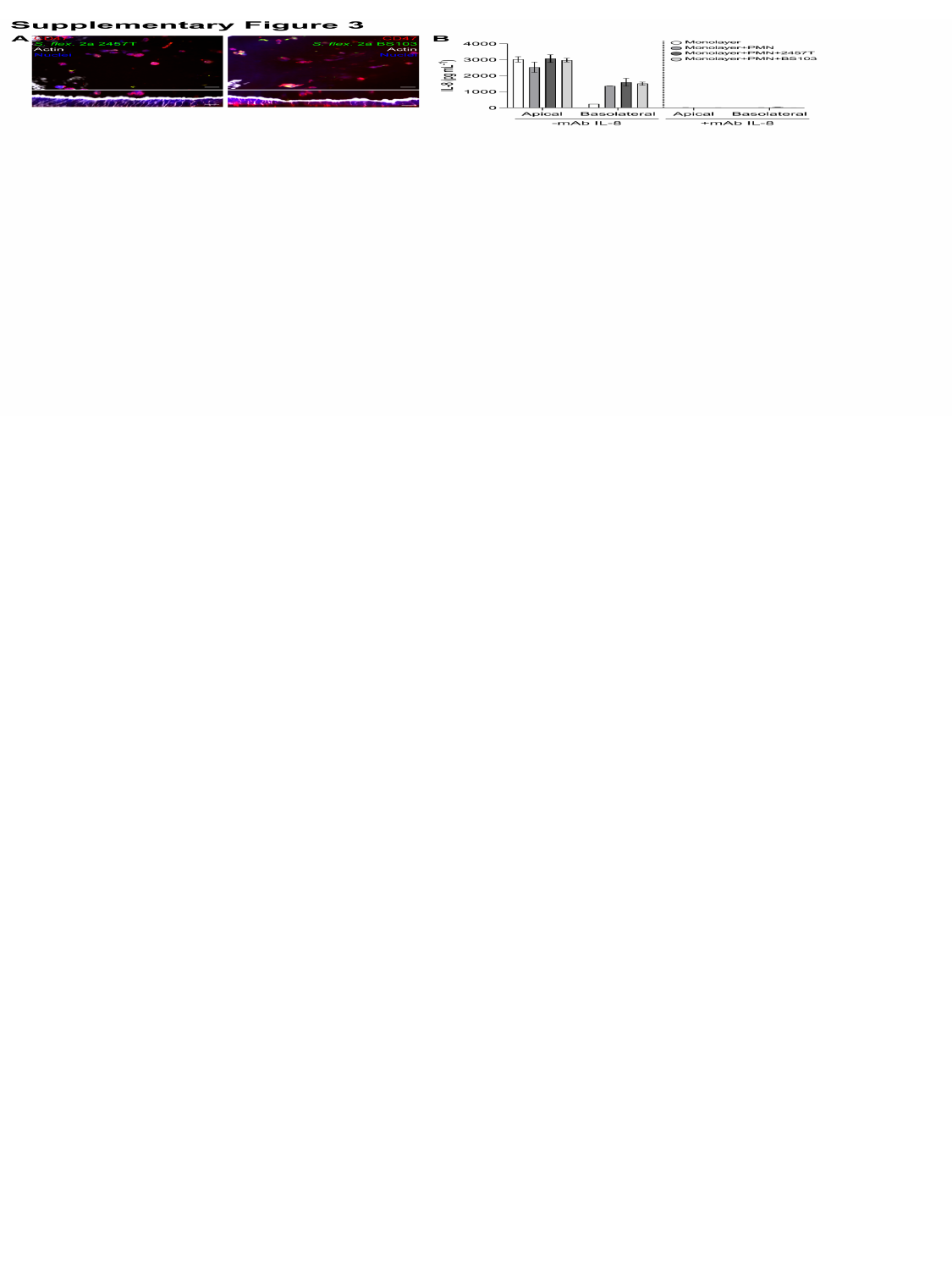

### Supplemental Figure 4

#### Slide 1
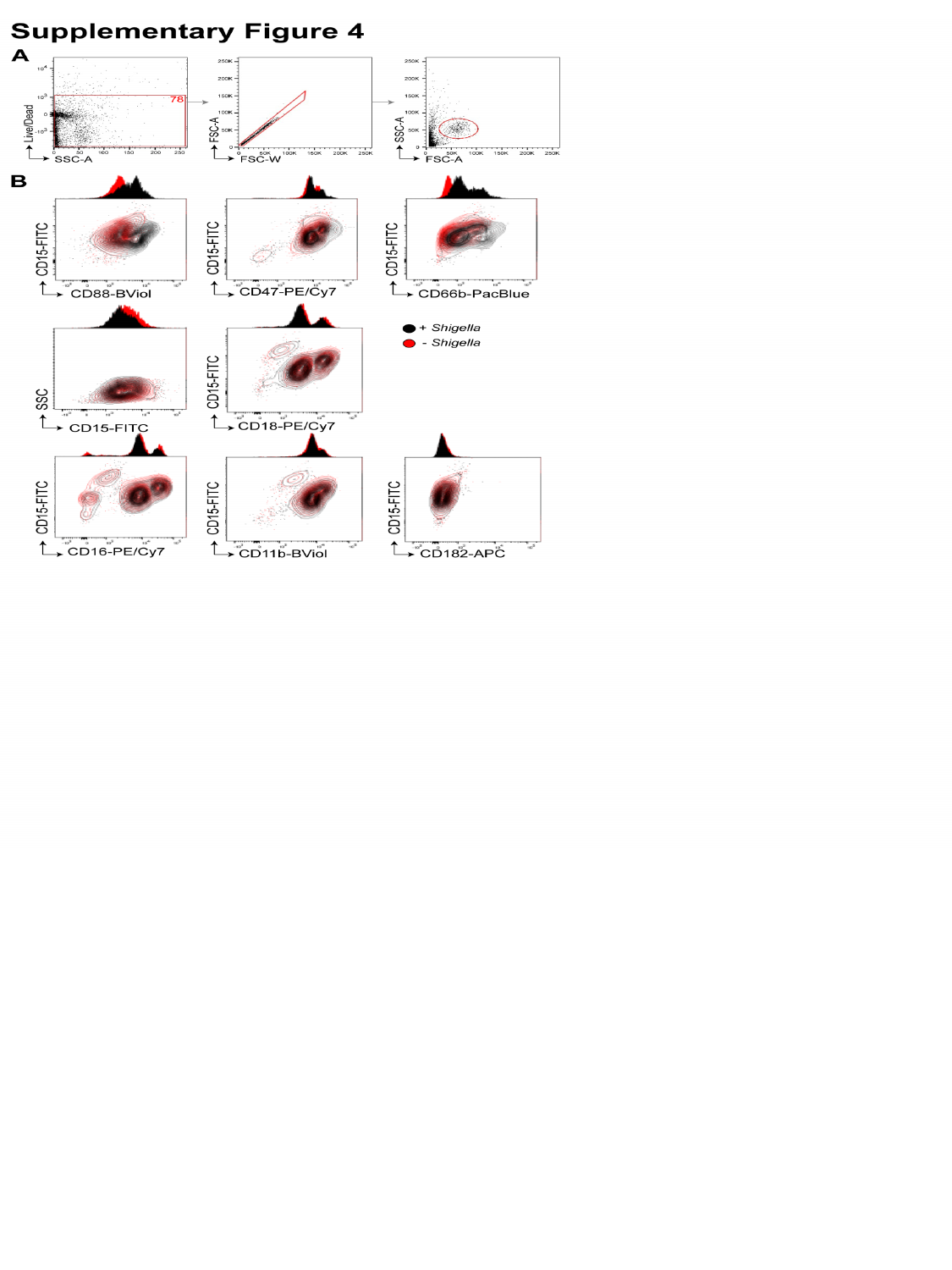
